# Deep Mutational Scanning Reveals EGFR Mutations Conferring Resistance to the 4th-generation EGFR tyrosine kinase inhibitor BLU-945

**DOI:** 10.1101/2025.03.31.646429

**Authors:** Yueyang Wang, Yuan Hao, Michela Ranieri, Michelle Hollenberg, Alfonso Lopez, Xavier T.R. Moore, Fiona Sherman, Jiehui Deng, Kwok-Kin Wong, Elaine Shum, John T. Poirier

**Affiliations:** Vilcek Institute of Graduate Biomedical Sciences, NYU Grossman School of Medicine; Laura and Isaac Perlmutter Cancer Center, NYU Langone Health

## Abstract

**Introduction:** Osimertinib, a covalent third-generation EGFR tyrosine kinase inhibitor (TKI) is the first-line standard of care for EGFR L858R and ex19del lung adenocarcinoma; however, tumors frequently acquire resistance through second-site mutations. Fourth-generation inhibitors designed to overcome common second-site resistance liabilities are in clinical development.

**Methods:** We performed deep mutational scanning (DMS) of the EGFR kinase domain in the context of an EGFR L858R driver mutation by transducing Ba/F3 cells with a saturation library of ∼17,000 EGFR L858R kinase domain variants. Ba/F3 cells expressing the DMS library were exposed to either osimertinib or BLU-945 to select for escape mutations.

**Results:** L718X mutations were enriched across all conditions as well as mutations private to BLU-945 treated samples including K714R, K716T, L718V, T725M, K728E, K754E/N, N771S/T, T783I, Q791L/K, G863S, S895N, K929I, and M971L. *In silico* pairwise comparisons of resistance profiles between each single agent condition suggested that combination treatment with osimertinib and BLU-945 would effectively suppress orthogonal resistance mechanisms, apart from L718X. A secondary DMS screen with osimertinib and BLU-945 in combination exclusively enriched for L718X mutations. L718X mutations were present in two patients treated with BLU-945 at our institution. One patient with both EGFR L858R and L718Q mutations prior to treatment was noted to have early progression. A second patient with *EGFR* L858R, T790M, and C797S at the time of enrollment acquired an L718V mutation at progression.

**Conclusions:** This study underscores the utility of comprehensive resistance profiles of single compounds, which can be used to predict the emergence of clinical resistance mutations and to devise combination treatments designed to suppress clonal escape.

## Introduction

Lung adenocarcinoma (LUAD) accounts for approximately 85% of primary lung cancer^1^, with most patients presenting with advanced or metastatic disease^2, 3^. Molecular profiling of LUAD has identified driver mutations in over 80% of cases^3^, with mutations in epidermal growth factor receptor (EGFR) among the most common^4^. Incidence in patients with Asian ancestry is 40-55%, notably higher than the 5-15% frequency observed in patients with Caucasian ancestry^5^. The majority of EGFR mutations occur in the kinase domain and manifest as short 3–7 amino acid deletions in exon 19 (ex19del) or as an L858R missense mutation encoded in exon 21^6, 7^. Less common mutations such as G719X and small insertions in exon 20 account for the remainder^6^. These mutations cause constitutive activation of EGFR, leading to unchecked cell proliferation, resistance to apoptosis and the promotion of metastasis^8^.

Targeting mutant EGFR with small molecule tyrosine kinase inhibitors (TKIs) has proven to be an effective therapeutic strategy in EGFR mutant LUAD, leading to multiple FDA approvals^9-12^. The first-generation TKIs gefitinib and erlotinib target the ATP-binding pocket of the kinase domain, effectively inhibiting downstream signaling^13^. Both gefitinib and erlotinib demonstrated significant improvement in progression-free survival (PFS) compared to platinum doublet chemotherapy in patients with tumors harboring EGFR L858R and ex19del alterations^11^. Despite initial clinical responses, acquired resistance frequently developed within one year^14^. A dominant mechanism of resistance to first-generation EGFR TKIs is the T790M Gatekeeper mutation, which abolishes TKI binding while retaining kinase activity^15, 16^.

One current standard of care for treatment-naïve EGFR mutant LUAD is osimertinib, a third-generation TKI, that was developed to overcome the T790M Gatekeeper mutation^17^. Osimertinib covalently target C797, inhibiting EGFR ex19del and L858R irrespective of the presence of T790M while sparing wild type EGFR^18^. Despite initial efficacy, 7-26% of patients develop resistance to osimertinib by acquiring additional second site mutations that are enriched in an allele specific manner^19, 20^. EGFR ex19del typically escapes osimertinib through L718Q/V mutations while L858R tends to escape *via* C797S, which destroys the cystine nucleophile required for covalent binding of osimertinib^21^.

In C797S-trans T790M mutations, where C797S and T790M are encoded on different alleles, combinations of first- and third-generation TKIs have shown some efficacy in preclinical models^18^. Conversely, in C797S-cis T790M mutations, where C797S and T790M are on the same allele, combination of first-generation TKIs with osimertinib are ineffective^18^. These data suggest that suppression of allelic diversity within cells or tumor cell populations may be an important consideration in the development of novel EGFR TKIs.

Fourth-generation ATP competitive EGFR TKIs are designed to retain efficacy in the context of C797S^22-24^. mutations; however, the landscape of potential second-site mutation liabilities associated with these inhibitors remains unknown^25, 26^. BLU-945 is an investigational fourth-generation EGFR TKI designed to target EGFR T790M and EGFR T790M/C797S mutations that confer resistance to first-, second- and third-generation TKIs^27^. Unlike osimertinib which binds covalently to the C797 residue, the scaffold of BLU-945 allows it to adopt a conformation that fits within the ATP binding pocket of EGFR in the presence of the C797S mutation while sparing wild-type EGFR^27^. BLU-945 has been evaluated clinically as a monotherapy and in combination with osimertinib in previously treated patients with EGFR driven LUAD^28^.

In this study, we aimed to identify potential mutational resistance mechanisms to BLU-945 using deep mutational scanning to assess the functional impact of thousands of EGFR mutations in parallel. We screened a DMS library of mutant cDNAs covering the EGFR L858R kinase domain to saturation in the context of a panel of EGFR TKIs to systematically profile all possible amino acid substitutions and their downstream effects.

## Methods

### Cell culture

HEK293T cells were cultured in high-glucose DMEM (Gibco #11995073) containing L-glutamine and sodium pyruvate, supplemented with 1% Penicillin-Streptomycin (Gibco #15140-122) and 10% Foundation™ fetal bovine serum (Gemini Bio-Products). Parental Ba/F3 cells were cultured in RPMI 1640 medium (Gibco #11875085) supplemented with 1% Penicillin-Streptomycin (Gibco #15140-122), 5% Foundation™ fetal bovine serum (Gemini Bio-Products), and 10 ng/mL recombinant mouse interleukin-3 (BioLegend #575504). Both HEK293T and Ba/F3 parental cell lines were obtained from the American Typed Culture Collection. All cell lines were maintained at 37°C in a humidified incubator with 5% CO_2_.

### Plasmids

The plasmid pHAGE-EGFR-L858R was a gift from Gordon Mills & Kenneth Scott (Addgene plasmid #116276; http://n2t.net/addgene:116276; RRID:Addgene_116276). pHAGE-EGFR-L858R was used as the backbone for subsequent deep mutational scanning (DMS) experiments. Plasmids were transformed and propagated in NEB Stable Competent *E*.*coli (*New England Biolabs #C3040H). To prepare the plasmid template for DMS library construction, the plasmid was linearized using the restriction enzyme BSU36I (New England Biolabs #R0524S), which recognizes distinct sequences flanking the region of interest. The digestion reaction was prepared in a final volume of 50 µL, consisting of 1 µg of plasmid DNA, 5 µL of 10× rCutSmart Buffer (New England Biolabs #B6004S), 1 µL of BSU36I, and nuclease-free water (New England Biolabs #B1500S) to bring the reaction to final volume. The mixture was incubated at 37°C for 2 hours to ensure complete digestion. Following digestion, the reaction products were resolved on a 0.7% agarose gel to confirm successful cleavage. The linear fragment encompassing the EGFR kinase domain was excised and purified using a gel extraction kit.

### Construction of EGFR L858R DMS library

To introduce mutations into the EGFR L858R kinase domain, we employed separate forward and reverse mutagenesis PCRs. Forward and reverse mutagenic primer were designed using the CodonTilingPrimers python script with minimum length set to 32 bp and synthesized as two separate oligonucleotide pools (Integrated DNA Technologies)^29^. In the forward reaction, 50 ng of linearized EGFR L858R template was combined with Q5 Master Mix (2×, New England Biolabs #M0491S), a 1 µL of a 10 µM forward oligo primer pool, and 1 µL of a 10 µM fixed reverse primer (5’-GCTCCAATAAATTCACTGCTTTG-3’) in a total volume of 50 µL. The PCR protocol was carried out for 7 cycles using standard Q5 cycling conditions, and the resultant product was purified with a PCR cleanup kit (Qiagen #28506). The reverse mutagenesis PCR was set up similarly using a fixed forward primer (5’-GCATACAGTGCCACCCAGAG-3’). The PCR protocol was also carried out for 7 cycles using standard Q5 cycling conditions, and the resultant product was purified with the PCR cleanup kit (Qiagen 28506). Each of the purified forward and reverse products was then treated with Exonuclease I (NEB M0293S) to remove any residual mutagenic primers. DNA concentrations were adjusted to approximately 140 ng/µL, and each product was further diluted by adding 60 µL of nuclease-free water to reach a final volume of 90 µL.

The forward and reverse products were joined to produce the full-length, mutated insert. Each 30 µL joining PCR reaction consisted of 15 µL of Q5 Master Mix (2×), 4 µL of the diluted forward PCR product, 4 µL of the diluted reverse PCR product, 1 µL of a 10 µM fixed forward primer, 1 µL of 10 µM fixed reverse primer, and 5 µL of nuclease-free water. Six parallel Joining PCRs were carried out to ensure sufficient product yield, using the same Q5 cycling conditions for 20 cycles. The products were then purified by PCR cleanup.

The mutagenized kinase domain was cloned into pHAGE-EGFR-L858R using restriction enzyme digestion and ligation. 1.6 µg of pHAGE-EGFR-L858R and 1 µg of the insert were digested with BSU36I and cleaned up by gel extraction. The digestion products were then ligated with T4 DNA ligase (New England Biolabs #M0202S) in a 20 µL reaction containing 166 ng of vector DNA, 100 ng of insert DNA, and 1 µL of T4 DNA ligase. The reaction was incubated at 16 °C overnight. A portion of the ligation mixture was tested by chemical transformation to confirm successful ligation prior to proceeding with large-scale transformation.

Electrotransformation was performed following the Endura™ Competent Cells (COMCEL-002) protocol. Briefly, 25 µL of electrocompetent cells were mixed with up to 5 µL of ligated DNA (final DNA concentration ∼55 ng/µL) in each electroporation cuvette. The SOC buffer was warmed to 37°C, and 5 mL culture tubes were prepared in advance. After electroporation, 970 µL of prewarmed SOC buffer were immediately added to each cuvette, and the cells were transferred to 5 mL tubes and recovered at 37°C for 1 hour with shaking at 250 rpm.

For library expansion, 1 mL of the recovered culture was plated on each Square BioAssay LB agar plate. To determine transformation efficiency by titration, ten-fold serial dilutions were prepared. Plates were incubated overnight at 37°C, and the resulting colony counts were used to calculate transformation efficiencies and ensure sufficient library complexity.

After overnight incubation at 37°C, the colonies on the large LB plate were harvested by gently scraping the plate surface with an appropriate volume of LB medium. The collected cells were transferred to a fresh tube, and plasmid DNA was extracted using a standard midi-prep protocol. This pooled plasmid DNA represented the final EGFR L858R DMS library. Individual colonies from the titration plates were selected for Sanger sequencing to determine the presence and frequency of intended mutations.

### Lentivirus production

On Day 0, HEK293T cells were seeded in a 15 cm dish to reach ∼70% confluence on Day 1. On Day 1, cells were transfected with 6 µg total DNA (EGFR L858R DMS library plasmid 3 µg, pSPAX 2 µg, pMD2.G 1 µg) in 200 µL Opti-MEM (Gibco #31985062), using a 2:1 ratio of linear polyethyleneimine (PEI) (Polysciences #23966) to total DNA. The DNA–PEI mixture was incubated at room temperature for 10 minutes before adding dropwise to the cells. On Day 2, the medium was refreshed. On Day 4, viral supernatants were collected, spun at 300 rcf for 5 minutes, and filtered through a 0.45 µm filter.

### Genomic DNA extraction

Genomic DNA (gDNA) was extracted from ∼1.5 × 10^7^ cells using salt precipitation^30^. Cells were resuspended in 200 µL of DPBS in a 15 mL tube. Lysis was achieved by adding 3 mL of lysis buffer (10 mM Tris, 100 mM EDTA, 2% SDS, pH 8.0) containing RNase A (10 mg/mL), followed by vortexing and incubation at 37°C for 5 minutes, then cooling on ice for 3 minutes. Protein contaminants were removed by adding 1 mL of protein precipitation solution (7.5 M ammonium acetate), vortexing, and centrifuging at 2,000 × g for 10 minutes.

The supernatant was transferred to a new tube containing 3 mL of isopropanol to precipitate DNA, and the mixture was gently inverted 50 times. DNA was pelleted by centrifugation at 2,000 × g for 3 minutes, washed with 3 mL of 70% ethanol, and air-dried for 5–10 minutes. The pellet was resuspended in 400 µL of hydration buffer (10 mM Tris, pH 8.0), incubated at 65°C for 1 hour, and hydrated overnight at room temperature with shaking at 600 rpm.

### Barcoded Subamplicon Sequencing

cDNAs were sequenced using a two-step, barcoded subamplicon PCR strategy based on the method described as previously described^29, 31^. In the first step (PCR1), the EGFR kinase domain was divided into two equal subamplicons, and each half was amplified in a separate reaction. Amplicon 1 used primers 5’-CTCTTGAGGATCTTGAAGGAAACTGAA-3’ and 5’-CTTGACATGCTGCGGTGTTTT-3’. Amplicon 2 used primers 5’-CTTGACATGCTGCGGTGTTTT-3’ and 5’-ATGCATTCTTTCATCCCCCTGAATGAC-3’. Approximately 5 µg of the DMS library DNA was split between these two reactions, each containing primers encoding 8 degenerate nucleotides to introduce unique molecular identifiers (UMIs). Following amplification, PCR1 products were purified by isopropanol precipitation to remove residual primers and short fragments, and the concentration of each purified product was determined using the PicoGreen method (Invitrogen #P11496).

For the second PCR step (PCR2), 500 fg of the purified PCR1 product was used as the template for each subamplicon. This PCR step incorporated Illumina sequencing adapters and sample-specific indexes for multiplexed sequencing. The resulting indexed libraries were then purified using SPRI beads.

Libraries were pooled at equimolar concentrations and sequenced on an Illumina NovaSeq SP (500 cycles) platform using paired end reads configured to capture the UMI, barcode, and subamplicon regions.

### Western blotting

Cells were lysed in RIPA buffer (Thermo Fisher #89900) supplemented with 1× Halt protease inhibitor cocktail (Thermo Fisher #78437) and phosphatase inhibitor cocktail (Thermo Fisher #78420). Total protein concentrations were determined using the BCA assay (Pierce). For each sample, 25 µg of total protein was mixed with 6× Laemmli SDS sample buffer (VWR, AAJ61337-AC) and heated at 95°C for 5 minutes. Samples were then separated on 4–12% gradient Bis-Tris gel and transferred onto PVDF membranes.

Membranes were blocked in 5% nonfat milk or 3% BSA (for phospho-specific detection) in TBS-T (20 mM Tris-HCl, 150 mM NaCl, 0.1% Tween-20) for 1 hour at room temperature. The blots were then incubated overnight at 4°C with primary antibodies against total EGFR (Cell Signaling Technology #2232S), phospho-EGFR (CST #3777S), total ERK (CST #9102S), phospho-ERK (CST #9101S), and cyclophilin B (CST #43603S), following the manufacturers’ recommended dilutions. After three washes in TBS-T, membranes were incubated with the HRP-conjugated secondary antibodies (Invitrogen #31460) at 1:10,000 for 1 hour at room temperature. Following three additional TBS-T washes, protein bands were visualized using the chemiluminescence substrate (Thermo Fisher #34577) and detected by the Invitrogen iBright imaging system.

### Cell viability assays

Ba/F3 cells were counted, adjusted to 20,000 cells/mL, and 25 µL (500 cells) was seeded into each well of a half-area 96-well plate (Corning #3688). Drug stocks were prepared at 2× concentrations using a 4-fold serial dilution (100 nM down to 0.0244 nM) in medium containing DMSO. Each drug concentration (including the DMSO-only control) was tested in triplicate by dispensing 25 µL of the 2× dilution into separate wells, yielding a final well volume of 50 µL at the 1× drug concentration. Plates were incubated at 37°C in 5% CO_2_ for 72 hours, after which 25 µL of CellTiter-Glo 2.0 reagent (Promega #G9242) was added per well. Following a 10-minute incubation at room temperature with gentle shaking, luminescence endpoint was measured on a microplate reader. Viability values were normalized to the DMSO control and used to generate dose–response curves for IC_50_ determination using GraphPad Prism (Dotmatics).

### Circulating tumor DNA sequencing

Peripheral blood was drawn at the NYU Perlmutter Cancer Center with the approval of the institutional review board (NYU IRB#S18-00980). Blood was collected in Streck Cell-Free DNA blood collection tubes and samples were mixed gently per manufacturer instructions, transported at 4°C, processed to aliquot serum, and subsequently stored at -80°C. Circulating tumor DNA sequencing was performed using the FoundationOne® Monitor (F1M) assay (Foundation Medicine, Inc.), a tissue-naïve method designed to monitor therapy response by quantifying ctDNA tumor fraction and variant allele frequency (VAF).

## Results

### Deep Mutational Scanning of the EGFR Kinase Domain Recovers Known Osimertinib Resistance Mutations

We generated a DMS library encompassing the EGFR L858R kinase domain encoded by a lentiviral transfer vector that co-expresses green fluorescent protein (GFP) *via* internal ribosomal entry site (IRES) **(Fig. 1A**). The library was generated using a mutagenic PCR approach, which relies on oligonucleotide primer pools encoding degenerate codons^29, 32, 33^. Each codon in the 804 bp target region can be mutated to one of 63 possible codons, including stop codons, resulting in a library of 16,884 potential variants (268 amino acids × 63 possible codons). The library consists of over 1×10^7^ primary clones, which corresponds to >500× representation over the theoretical number of variants. Sanger sequencing of individual clones indicated an average of 1.14 mutations per cDNA (**Fig. 1B**). The plasmid library was sequenced using barcoded error correction and showed negligible mutational bias in transitions and transversions (**Fig. 1C**). 16,882 unique variants were recovered, corresponding to >99.9% saturation with >99% of variants observed >10 times (**Fig. 1D)**. The abundance of codon variants was approximately normally distributed with a mean of 168 unique observations (**Fig. 1E**). The EGFR L858R DMS library was packaged in VSV-g pseudotyped lentivirus at large scale and concentrated by centrifugation. The titer of the lentiviral library was 2×10^7^ transduction units (TU) as determined by detection of GFP expression by flow cytometry. The total TU corresponded to ∼1,185× representation.

**Figure 1.**
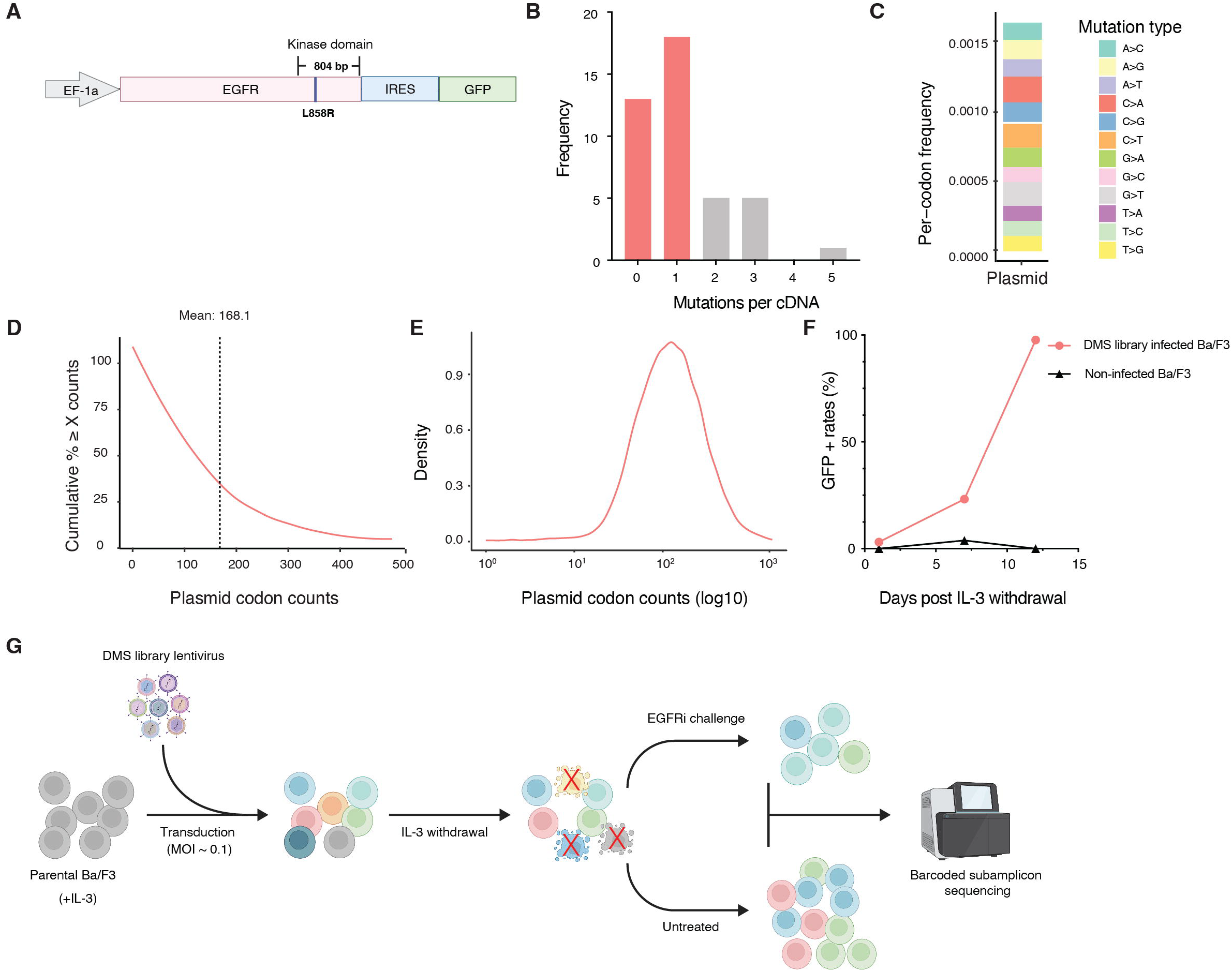
Deep mutational scanning of the EGFR L858R kinase domain in Ba/F3 cells. **A)** Map of pHAGE-EGFR-L858R with kinase domain annotated. **B)** Frequency of mutations count per cDNA **C)** Per-codon frequency of the indicated mutation types **D)** Cumulative distribution of read counts for all expected variants in the library. **E)** Density plot of plasmid library codon counts **F)** Enrichment of library transduced Ba/F3 cells after IL-3 withdrawal. **G)** Positive selection screening scheme.

We conducted all screens in Ba/F3 cells, a murine pro-B cell line dependent on extracellular interleukin-3 (IL-3) for survival. Upon transduction with potent oncogenes, these cells become dependent on the introduced oncogene for mitogenic signaling^34^. Ba/F3 cells were transduced at a low multiplicity of infection (MOI ∼0.1) to bias toward single integrations while achieving >1,000-fold representation of the library^29, 31^. After allowing 7 days for provirus integration and EGFR L858R mutant protein expression, IL-3 was withdrawn from the culture media. This established a selective pressure in which Ba/F3 cell survival depended on the signaling activity of transduced EGFR mutants^35^. Cells expressing viable EGFR variants were expected to survive without IL-3, while those harboring non-functional or deleterious mutations, or that were not transduced, were outcompeted and depleted from the population. After 14 days of IL-3 withdrawal, over 95% of the surviving Ba/F3 cells were GFP positive, consistent with positive selection of the transduced pool (**Fig. 1F)** ^31, 36, 37^.

To test whether known resistance mutations could be recovered from the Ba/F3 DMS pool, we exposed the EGFR L858R DMS library-expressing Ba/F3 cells to osimertinib at 100nM (>EC_90_ of osimertinib) over a 14-day period with regular replenishment of drug to maintain consistent pressure (**Fig. 1G**). Endpoint samples were harvested, gDNA was extracted, and deep sequencing with error correction was performed as previously described^31^. Independent biological replicates demonstrated high concordance in the enriched variant profiles, supporting the reproducibility of the results (**Supplementary Fig. 1)**. Among the osimertinib-treated samples, L718X mutations were enriched compared to control. Since L718X is the dominant resistance mutation to osimertinib in the context of L858R mutation, we concluded that the screening approach was valid.

### Deep Mutational Scanning Reveals EGFR Mutations Conferring Resistance to BLU-945

We employed the same screening workflow in the context of BLU-945 at 100 nM (>EC_90_) for 14 days with regular drug replenishment. Endpoint samples were harvested, gDNA was extracted, and deep sequencing was performed as above. In contrast to the osimertinib screen, we identified numerous potential resistance mutations for BLU-945 (**Fig. 2A**). We identified mutations clustered in the hinge region amino acids Q791 and M793, which form hydrogen bonds with the adenosine ring of ATP^38^. Q791L/K and M793X mutations are supported as likely resistance mutations by structural evidence of a previously published co-crystal structure of the BLU-945 analog ‘compound 24’ and EGFR kinase domain (PDB:8D76)^27^. The amino-naphthyridine NH group of BLU-945 forms a hydrogen bond to the backbone carbonyl of Q791 while the naphthyridine N2 forms a hydrogen bond with the backbone NH of M793, stabilizing the inhibitor-kinase complex. We conclude that the results of the screen are consistent with the binding mode of BLU-945.

**Figure 2.**
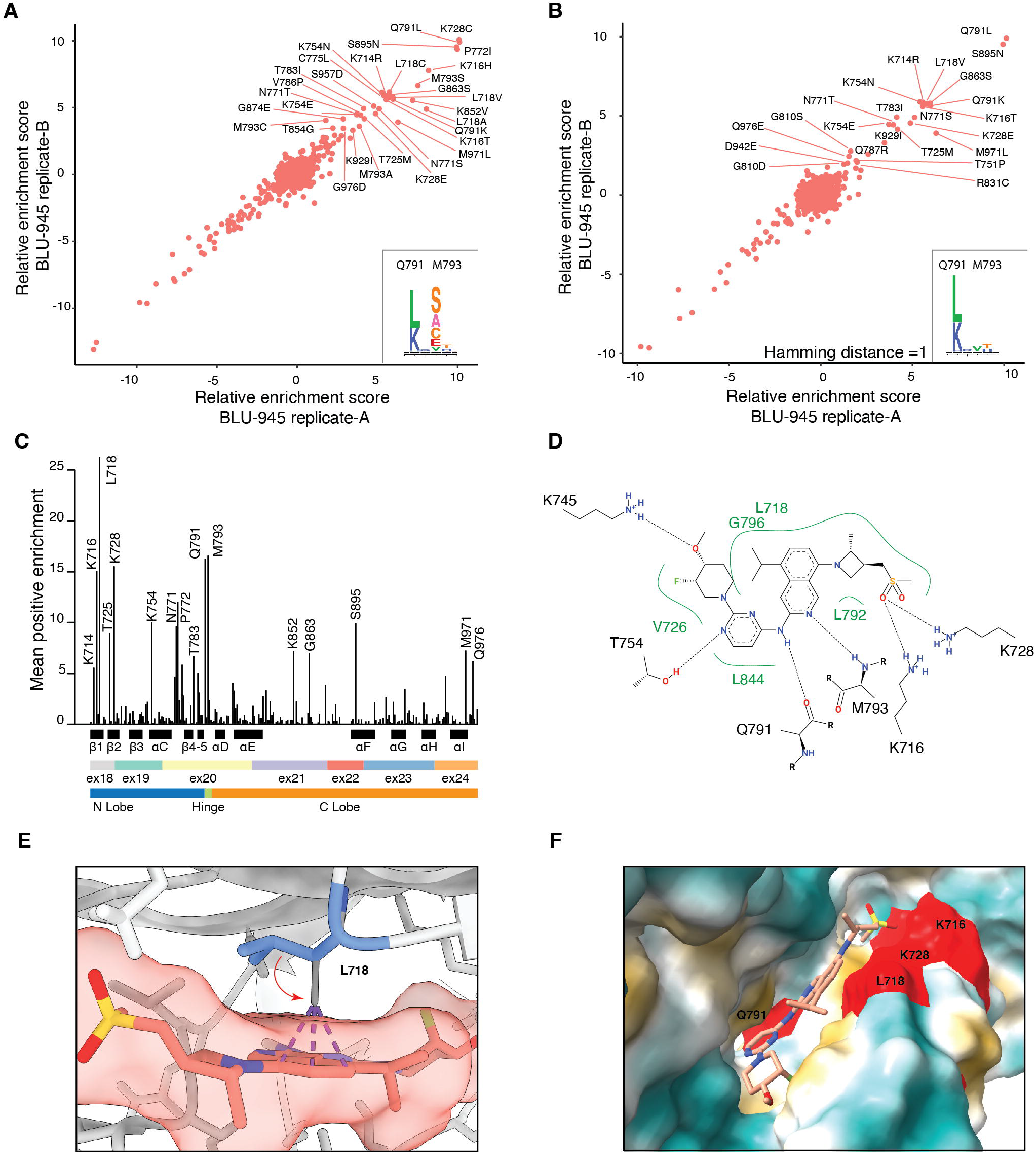
Identification of resistance mutations for osimertinib and BLU-945. **A)** Scatter plot of enrichment scores after BLU-945 selection. Inset: hinge mutations involved in APT-binding. **B)** Scatter plot of enrichment scores after BLU-945 selection, Hamming distance of 1. Inset: hinge mutations present after filtering. **C)** Enrichment of BLU-945 resistance mutations relative to the EGFR kinase domain structure. **D)** Poseview diagram of BLU-945, illustrating key contact residues. **E)** Illustration of predicted clashes between BLU-945, and a likely rotamer of L718V. **F)** Resistance mutations that may disrupt key hydrogen bonds. Frequently mutated residues are highlighted in red.

DMS screens explore all potential mutations at each amino acid position that are compatible with sustained EGFR L858R signaling, including complex codon changes that are less likely to occur in patients. We focused on amino acid changes that could arise from a single nucleotide substitution (Hamming distance = 1) from the wild-type codon. After filtering, Q791L/K mutations remain while M793X are removed, suggesting that mutations to M793 would be an unlikely escape mechanism (**Fig. 2B**). We identified the following mutations as resistance mutations that can arise through single base mutations: K714R, K716T, L718V, T725M, K728E, K754E/N, N771S/T, T783I, Q791L/K, G863S, S895N, K929I, and M971L.Several of these mutations have been observed clinically, such as K714R^39^, L718V^40^, T725M^41^, K728E^42^, T751P^43, 44^, K754E/N^45, 46^, and N771S^47^.

When viewing BLU-945 mutations in the context of kinase domain structure, several patterns become apparent (**Fig. 2C**). First, mutations tend to be clustered in the N lobe and hinge regions of the kinase domain, which play a comparatively larger role in the catalytic activity of EGFR than the C lobe. Second, we observed frequent mutations in lysine residues in the β1-3 strands of the N lobe. Third, we see a cluster of mutations in the region between the αC-helix and β4-5 strands involving N771 and P772, two residues involved in ex20ins variants^48^. We suspect that these point mutations in this region might distort P-loop and αC-helix conformation in a manner that indirectly affects BLU-945 binding.

Many of the mutations highlighted here involve contact residues that are affinity determinants of BLU-945 (**Fig. 2D**). Notably, Q791 and M793 hinge contact points were present in compound 7, an early lead compound in the BLU-945 discovery campaign, while other mutations such as K716T and K728E involve residues engaged by the sulfone group characteristic of compound 24 and eventually BLU-945, developed late in the medicinal chemistry program. The large number of potential mutational escape routes for BLU-945 relative to osimertinib may be due to a greater number of required binding affinity determinants.

Not observed in the screen were mutations to the back pocket residue K745, which forms a hydrogen bond with the piperidinol ring of BLU-945^27^. Molecular interactions with K745 are a common and distinct feature of reversible, fourth-generation ATP competitive EGFR kinase inhibitors^38^. We suspect that K745 mutations are not observed since this residue is the conserved “catalytic lysine” which is necessary to form a salt bridge with E762 in the kinase active state that is essential for phosphoryl transfer activity^49^.

In addition to mutations of key contact point residues, DMS identified L718V as a key resistance mutation to BLU-945, similar to the osimertinib screen. Structurally, L718 is positioned in the first β-strand (β1) of the N-lobe near the ATP-binding site. When leucine at position 718 is substituted by valine, the valine side chain at position 718 is predicted to rotate ∼180º into the ATP-binding pocket and clash with BLU-945, which could prevent the inhibitor from properly occupying the pocket (**Fig. 2E**). Overall, the key residues involved in BLU-945 binding that are liabilities for mutational escape tend to be clustered around the ATP-binding pocket occupied by BLU-945 (**Fig. 2F**).

### Functional validation of individual BLU-945 resistance mutations in multiple allelic contexts

To test whether any resistance mutations identified in the primary screen were sufficient to confer resistance to BLU-945, we chose to focus on the Q791L hinge mutation given its importance in the primary screen for BLU-945 and the relative ease of mutation at that site as well as L718Q, since this mutation is most frequently identified as a resistance mutation in patients with L858R mutations^27, 50^. We generated individual Ba/F3 cell lines stably expressing candidate resistance mutations in the context of L858R and del19 mutations (delE746-A750), which represent the most common EGFR-activating mutations in lung adenocarcinoma. Additionally, we introduced co-mutations of T790M and C797S to model clinically relevant resistance scenarios. All mutant constructs conferred growth factor independence in Ba/F3 cells apart from del19/L718Q, consistent with published reports in patients and mouse model systems suggesting that this co-mutation is non-functional^51, 52^.

We observed that Ba/F3 cells in L858R or del19 context, in conjunction with T790M/C797S mutations, displayed increased resistance to BLU-945 when either L718Q or Q791L mutations were present. The EC_50_ values for L858R/T790M/C797S and del19/T790M/C797S were 1.71 nM (1.26nM – 2.40nM 95%CI) and 7.81 nM (6.09nM – 9.96nM 95%CI) respectively; however, the EC_50_ was >100nM with the addition of L718Q or Q791L resistance mutations. (**Fig. 3A-D**).

**Figure 3.**
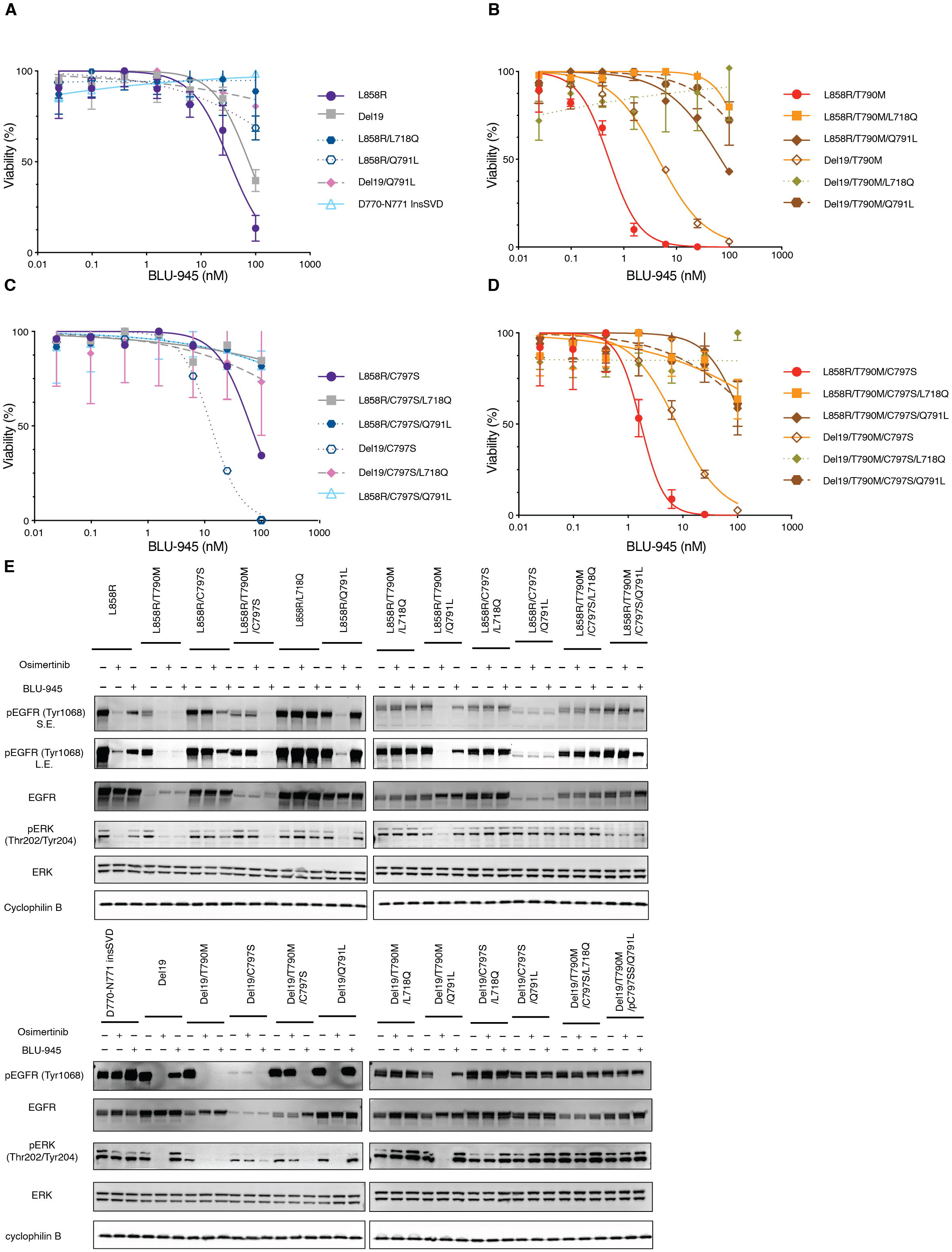
Testing of individual resistance mutations in the context of L858R, ex19del, T790M, and C797S. **A)** Viability dose-response curves showing the sensitivity of Ba/F3 cells expressing EGFR L858R or Del19 alone, as well as those harboring candidate resistance mutations (L718Q, Q791L), (D770-N771 InsSVD as a EGFR ex20ins control). **B)** Viability dose-response curves of Ba/F3 cells expressing EGFR L858R/T790M or Del19/T790M, with or without co-occurring L718Q or Q791L mutations. **C)** Viability doseresponse curves of Ba/F3 cells expressing EGFR L858R/C797S or Del19/C797S, with or without cooccurring L718Q or Q791L mutations. **D)** Viability dose-response curves of Ba/F3 cells harboring the triplet mutation combinations L858R/T790M/C797S or Del19/T790M/C797S, with or without L718Q or Q791L. **E)** Western blot analysis of Ba/F3 cells under the indicated conditions.

Western blot analysis revealed that the phosphorylation levels of EGFR in L858R/T790M/C797S and del19/T790M/C797S mutant cells were markedly reduced following treatment with 100nM BLU-945. However, no significant reduction in phosphorylation was observed in cells harboring L718Q or Q791L cDNAs, indicating that these second-site mutations confer resistance to BLU-945 by escaping EGFR inhibition (**Fig 3E**).

### Osimertinib co-treatment suppresses BLU-945 kinase domain resistance mutants apart from L718X

In the SYMPHONY trial, BLU-945 was tested in combination with osimertinib^28^. To explore the potential of overcoming resistance to BLU-945 through combination therapy, we tested whether the resistance profiles of individual compounds, in this case BLU-945 and osimertinib, could predict the resistance profile for the combination treatment. Pairwise comparison of single agent resistance profiles suggested that osimertinib could effectively inhibit most resistance mutations observed with BLU-945 monotherapy, with the notable exception of mutations at position L718 (**Fig. 4A**).

**Figure 4.**
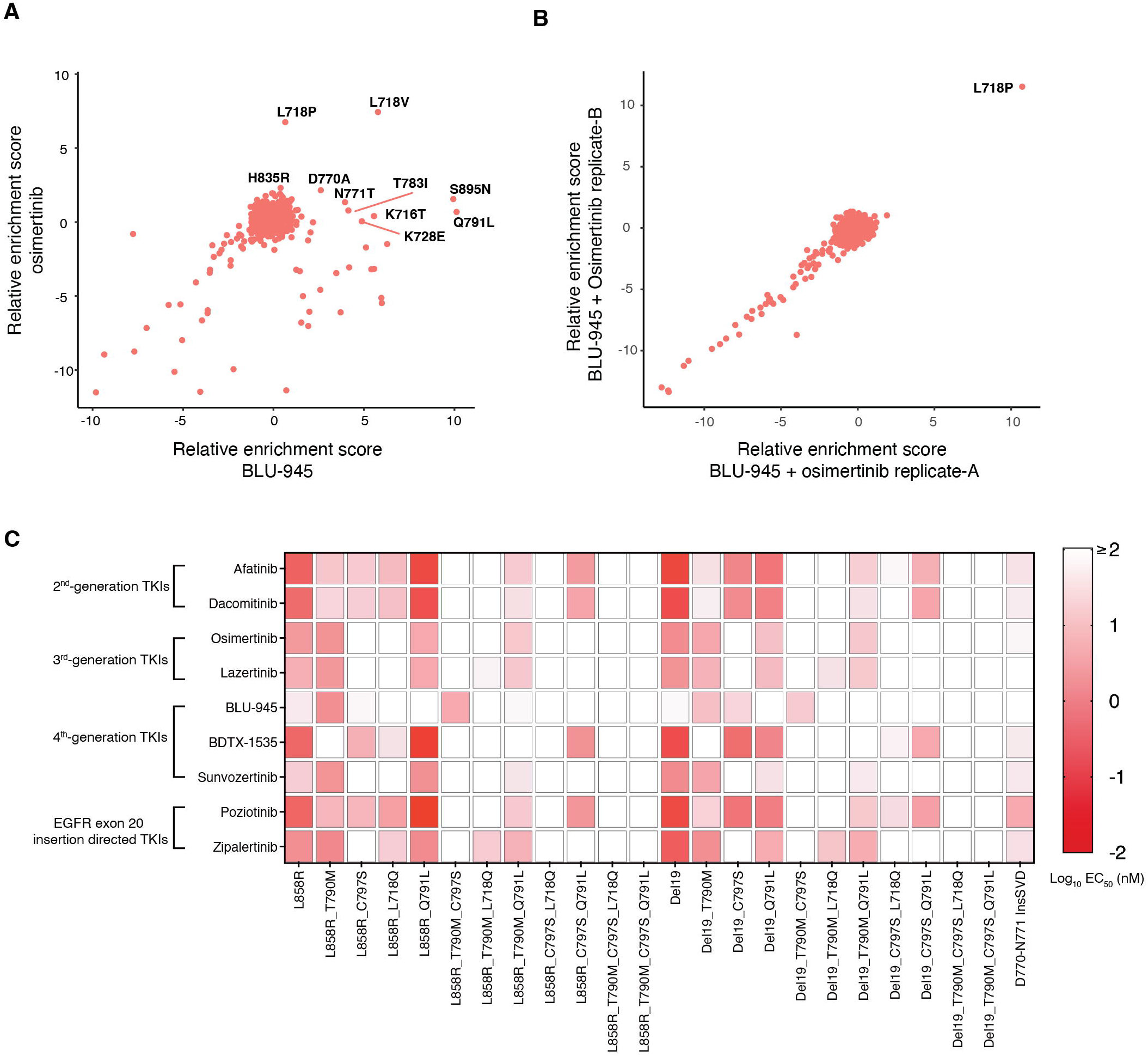
Prediction and experimental validation of EGFRi cross-resistance. **A)** Pairwise comparison of Osimertinib and BLU-945 resistance profiles. **B)** Scatter plot of enrichment scores after combination osimertinib and BLU-945 selection. **C)** Heatmap of Log_10_ EC_50_ values for 10 distinct inhibitors calculated from 7-point dose response curves.

To test our hypothesis, we performed a DMS experiment using BLU-945 and osimertinib in combination. The enrichment plot generated from these co-treatment endpoint samples indicated that resistance to both agents exclusively occurs at the L718 position where only proline (L718P) is enriched after filtering for Hamming distance >1 (**Fig. 4B**), illustrating a common resistance liability at this position.

While osimertinib and BLU-945 share a common resistance liability at position L718, we were encouraged that all other second-site mutations from the BLU-945 screen were effectively suppressed. We tested the resistance profiles of 9 distinct EGFR inhibitors to determine to what extent they shared resistance mechanisms with BLU-945 (**Fig. 4C**). Notably, BLU-945 suppressed T790M/C797S in both L858R and ex19del contexts. Mutations conferring resistance to BLU-945 such as Q791L were sensitive to all drugs tested in the context of either L858R or ex19del. However, rescue by osimertinib, sunvozertinib, lazertinib, or zipalertinib were all abolished by the C797S co-mutation. Examination of cross-resistance patterns of these drugs suggests that the combination of BLU-945 with lazertinib might be an attractive strategy based on coverage of both C797S and L718Q resistance mutations in contexts where the two mutations do not occur *in cis*.

### BLU-945 and osimertinib cross-resistant L718X mutations identified in two patients with primary resistance and on-treatment disease progression

Our institution participated in the first-in-human Phase I/II SYMPHONY clinical trial evaluating BLU-945, both as monotherapy and in combination with osimertinib, for patients with metastatic EGFR-mutant NSCLC. We identified two patients harboring L718X EGFR mutations.

Patient A (**Fig. 5A**) harbored an EGFR L858R mutation and was previously treated with osimertinib. The patient had a liquid biopsy obtained on day –659 which identified the presence of *EGFR* L858R and L718Q mutations. Baseline imaging on day 0 revealed two nodules in the right lung measuring 1.2 × 0.7 cm (middle lobe) and 2.7 × 2.2 cm (lower lobe). The patient started BLU-945 on day 12. Although initial follow-up imaging demonstrated stable disease, evaluations on day 37 and day 40 revealed disease progression with new bone metastases and spinal cord compression, necessitating surgical intervention. Post-surgery imaging on day 68 showed stable lung lesions, but further scans on day 121 demonstrated continued disease progression (+20% from nadir) along with new brain metastases. Additional progression in the spine was confirmed on day 177 and day 181.

**Figure 5.**
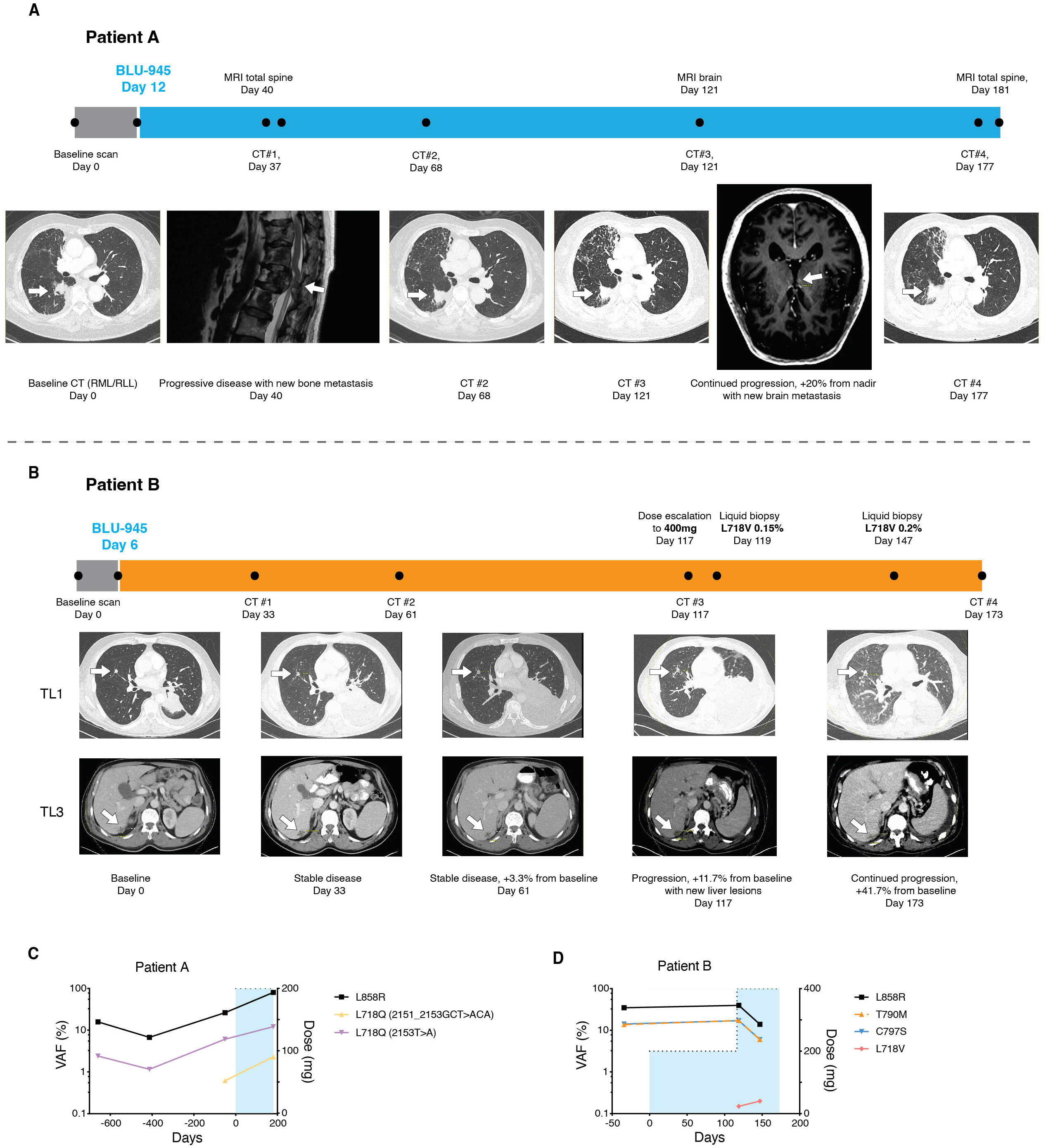
Clinical courses of two patients with L718X mutation. **A)** Clinical course of patient A. **B)** Clinical course of patient B. **C)** ctDNA sequencing results for patient A. Left axis is log_10_ variant allele frequency, right axis is BLU-945 dose. **D)** ctDNA sequencing results for patient B. Left axis is log_10_ variant allele frequency, right axis is BLU-945 dose.

Patient B (**Fig 5B**) presented with *EGFR* L858R and developed on-treatment resistance mutations (*EGFR* T790M and C797S) following multiple prior treatments including osimertinib. A pre-treatment liquid biopsy performed on day -34 revealed *EGFR* L858R (34.7% VAF), T790M (13.7% VAF), and C797S (14.2% VAF). Baseline imaging on day 0 showed a right middle lobe lung nodule measuring 1.0 × 0.8 cm (TL1) as well as a liver lesion measuring 1.5×1.1cm (TL3). BLU-945 therapy commenced on day 6. Follow-up imaging on day 33 demonstrated stable disease, with the sum of the target lesions decreasing by −1.7% overall from baseline. However, by day 61, the sum of target lesions increased by 3.3%. On day 117, imaging demonstrated progression due to new liver lesions, prompting a BLU-945 dose escalation to 400 mg.

On day 173, CT imaging indicated an overall growth of the target lesions by > 41%. Although the patient initially exhibited stable disease, therapy was ultimately discontinued due to rapid disease progression.

Four liquid biopsies were obtained from Patient A at day -659, -413, -51, and 180 relative to day 0 (**Fig. 5C**). During this interval, the *EGFR* L858R mutation rose from 15.8% to 80.0% VAF. Two distinct L718Q variants were also detected: L718Q (2151_2153GCT>ACA) was first measurable at 0.62% on day -51 and increased to 2.29% by day 180. L718Q (2153T>A) was detected at 2.4% on day -659, fluctuating thereafter and ultimately reaching 12.11% on day 180.

Two on-treatment liquid biopsies were obtained from Patient B on days 119 and 147. The first confirmed *EGFR* L718V (0.15% VAF) in addition to L858R (39.69% VAF), T790M (17.19% VAF), and C797S (17.01% VAF) (**Fig. 5D**). Liquid biopsy on day 147 coincided with an increase in L718V to 0.2% VAF, with reduced levels of L858R (13.9% VAF), T790M (6% VAF), and C797S (6% VAF).

## Discussion

In this study, we employed deep mutational scanning to systematically map amino acid substitutions in the EGFR L858R kinase domain that confer resistance to BLU-945, a fourth-generation EGFR inhibitor. We validated Q791L and L718Q mutations, which emerged as crucial resistance determinants in L858R and ex19del contexts irrespective of T790M or C797S co-mutations. L718X mutations are notable as a singular shared liability between BLU-945 and osimertinib, highlighting the challenge of covering all potential mutational escape routes even in the case of structurally distinct inhibitors.

DMS has the potential to provide comprehensive foreknowledge of on-target resistance liabilities not previously possible by methods such as long-term drug treatment or chemical mutagenesis. This approach can be integrated into early drug development pipelines to refine chemical scaffolds against known escape mutations. We show that forward genetics can complement structural information by highlighting constrained contact points such as M793 and K745, while minimizing the vulnerability at sites like Q791 and L718. These positions in the EGFR kinase domain may be more “mutationally pliable” than others, enabling multiple potential substitutions that preserve kinase function and reduce inhibitor efficacy. Defining these mutationally permissive hotspots could support the development of inhibitors with increased robustness to mutational mechanisms of acquired resistance.

Anticipating drug resistance mutations early in drug development could also inform rational design of combination or sequential regimens in the clinic. For example, if preclinical data predict that L718X and Q791L/K will arise under selective pressure from BLU-945, upfront strategies could include pairing BLU-945 with a complementary inhibitor with orthogonal resistance liabilities to diminish the potential for outgrowth of resistant clones. From the perspective of clinical trial design, comprehensive resistance profiles could be used to further stratify patients to enhance response rates, increasing the probability of successful trial. This approach would also respect ethical considerations by providing the necessary information to direct patients away from therapies that have reduced potential for efficacy.

Our study is not without limitations. Our screen focuses entirely on the EGFR kinase domain in the context of a single L858R mutant allele. Mutations outside the kinase domain or that require different allelic context would be missed. On target mutations represent only one facet of clinical resistance, whereas real-world tumors can escape *via* other genetic processes such as gene amplification, bypass signaling, or histological transformations^53^. The Ba/F3 system, while well-established for oncogene screening, may not fully capture the complexities of human lung adenocarcinoma. Screening at a single high-dose condition (>EC_90_) could potentially miss subtle resistance phenotypes that might emerge with lower drug concentrations. Finally, while our analysis tested candidate mutations in ex19del and L858R, our primary screen focused on the L858R context, whereas exon ex19del, ex20ins, or comparatively less common point mutations such as G719X or L861Q might yield distinct allele-specific resistance mutations.

Overall, these results illustrate the power of systematic mutational scanning to identify resistance liabilities for emerging TKIs. Future studies could build upon these findings by exploring combination strategies, building prospective surveillance of likely resistance mutations into early phase clinical trials, and designing novel TKIs that target common resistance escape routes.

## Notes

### Competing Interest Statement

Costs of this research were partially covered by Blueprint Medicines.

## References

1. Midha A, Dearden S, McCormack R. mutation incidence in non-small-cell lung cancer of adenocarcinoma histology: a systematic review and global map by ethnicity (mutMapII). Am J Cancer Res. 2015;5(9):2892–+. PubMed PMID: WOS:000364773000028.

2. Crino L, Weder W, van Meerbeeck J, Felip E, Group EGW. Early stage and locally advanced (non-metastatic) non-small-cell lung cancer: ESMO Clinical Practice Guidelines for diagnosis, treatment and follow-up. Ann Oncol. 2010;21 Suppl 5:v103–15. doi: 10.1093/annonc/mdq207. PubMed PMID: 20555058.

3. Dearden S, Stevens J, Wu YL, Blowers D. Mutation incidence and coincidence in non small-cell lung cancer: meta-analyses by ethnicity and histology (mutMap). Annals of Oncology. 2013;24(9):2371–6. doi: 10.1093/annonc/mdt205. PubMed PMID: WOS:000323963100026.

4. Smits AJJ, Kummer JA, Hinrichs JWJ, Herder GJM, Scheidel-Jacobse KC, Jiwa NM, Ruijter TEG, Nooijen PTGA, Looijen-Salamon MG, Ligtenberg MJL, Thunnissen FB, Heideman DAM, de Weger RA, Vink A. and mutations in lung carcinomas in the Dutch population: increased mutation frequency in malignant pleural effusion of lung adenocarcinoma. Cell Oncol. 2012;35(3):189–96. doi: 10.1007/s13402-012-0078-4. PubMed PMID: WOS:000306403400007.

5. Li KD, Yang MJ, Liang NX, Li SQ. Determining EGFR-TKI sensitivity of G719X and other uncommon EGFR mutations in non-small cell lung cancer: Perplexity and solution (Review). Oncol Rep. 2017;37(3):1347–58. doi: 10.3892/or.2017.5409. PubMed PMID: WOS:000396186600006.

6. Harrison PT, Vyse S, Huang PH. Rare epidermal growth factor receptor (EGFR) mutations in non-small cell lung cancer. Semin Cancer Biol. 2020;61:167–79. doi: 10.1016/j.semcancer.2019.09.015. PubMed PMID: WOS:000520017900017.

7. Eck MJ, Yun CH. Structural and mechanistic underpinnings of the differential drug sensitivity of EGFR mutations in non-small cell lung cancer. Biochim Biophys Acta. 2010;1804(3):559–66. Epub 20091222. doi: 10.1016/j.bbapap.2009.12.010. PubMed PMID: 20026433; PMCID: PMC2859716.

8. Uribe ML, Marrocco I, Yarden Y. EGFR in Cancer: Signaling Mechanisms, Drugs, and Acquired Resistance. Cancers (Basel). 2021;13(11). Epub 20210601. doi: 10.3390/cancers13112748. PubMed PMID: 34206026; PMCID: PMC8197917.

9. Lynch TJ, Bell DW, Sordella R, Gurubhagavatula S, Okimoto RA, Brannigan BW, Harris PL, Haserlat SM, Supko JG, Haluska FG, Louis DN, Christiani DC, Settleman J, Haber DA. Activating mutations in the epidermal growth factor receptor underlying responsiveness of non-small-cell lung cancer to gefitinib. N Engl J Med. 2004;350(21):2129–39. Epub 20040429. doi: 10.1056/NEJMoa040938. PubMed PMID: 15118073.

10. Paez JG, Janne PA, Lee JC, Tracy S, Greulich H, Gabriel S, Herman P, Kaye FJ, Lindeman N, Boggon TJ, Naoki K, Sasaki H, Fujii Y, Eck MJ, Sellers WR, Johnson BE, Meyerson M. EGFR mutations in lung cancer: correlation with clinical response to gefitinib therapy. Science. 2004;304(5676):1497–500. Epub 20040429. doi: 10.1126/science.1099314. PubMed PMID: 15118125.

11. Mok TS, Wu YL, Thongprasert S, Yang CH, Chu DT, Saijo N, Sunpaweravong P, Han B, Margono B, Ichinose Y, Nishiwaki Y, Ohe Y, Yang JJ, Chewaskulyong B, Jiang H, Duffield EL, Watkins CL, Armour AA, Fukuoka M. Gefitinib or carboplatin-paclitaxel in pulmonary adenocarcinoma. N Engl J Med. 2009;361(10):947–57. Epub 20090819. doi: 10.1056/NEJMoa0810699. PubMed PMID: 19692680.

12. Soria JC, Ohe Y, Vansteenkiste J, Reungwetwattana T, Chewaskulyong B, Lee KH, Dechaphunkul A, Imamura F, Nogami N, Kurata T, Okamoto I, Zhou C, Cho BC, Cheng Y, Cho EK, Voon PJ, Planchard D, Su WC, Gray JE, Lee SM, Hodge R, Marotti M, Rukazenkov Y, Ramalingam SS, Investigators F. Osimertinib in Untreated EGFR-Mutated Advanced Non-Small-Cell Lung Cancer. N Engl J Med. 2018;378(2):113–25. Epub 20171118. doi: 10.1056/NEJMoa1713137. PubMed PMID: 29151359.

13. El-Rayes BF, LoRusso PM. Targeting the epidermal growth factor receptor. Br J Cancer. 2004;91(3):418–24. doi: 10.1038/sj.bjc.6601921. PubMed PMID: 15238978; PMCID: PMC2409851.

14. Wu SG, Shih JY. Management of acquired resistance to EGFR TKI-targeted therapy in advanced non-small cell lung cancer. Mol Cancer. 2018;17. doi: ARTN 38 10.1186/s12943-018-0777-1. PubMed PMID: WOS:000425760100006.

15. Pao W, Miller VA, Politi KA, Riely GJ, Somwar R, Zakowski MF, Kris MG, Varmus H. Acquired resistance of lung adenocarcinomas to gefitinib or erlotinib is associated with a second mutation in the EGFR kinase domain. Plos Med. 2005;2(3):225–35. doi: ARTN e73 10.1371/journal.pmed.0020073. PubMed PMID: WOS:000228382500014.

16. Kobayashi S, Boggon TJ, Dayaram T, Janne PA, Kocher O, Meyerson M, Johnson BE, Eck MJ, Tenen DG, Halmos B. EGFR mutation and resistance of non-small-cell lung cancer to gefitinib. N Engl J Med. 2005;352(8):786–92. doi: 10.1056/NEJMoa044238. PubMed PMID: 15728811.

17. Mok TS, Wu YL, Ahn MJ, Garassino MC, Kim HR, Ramalingam SS, Shepherd FA, He Y, Akamatsu H, Theelen WS, Lee CK, Sebastian M, Templeton A, Mann H, Marotti M, Ghiorghiu S, Papadimitrakopoulou VA, Investigators A. Osimertinib or Platinum-Pemetrexed in EGFR T790M-Positive Lung Cancer. N Engl J Med. 2017;376(7):629–40. Epub 20161206. doi: 10.1056/NEJMoa1612674. PubMed PMID: 27959700; PMCID: PMC6762027.

18. Fu K, Xie FC, Wang F, Fu LW. Therapeutic strategies for EGFR-mutated non-small cell lung cancer patients with osimertinib resistance. J Hematol Oncol. 2022;15(1). doi: ARTN 173 10.1186/s13045-022-01391-4. PubMed PMID: WOS:000895918400002.

19. Leonetti A, Sharma S, Minari R, Perego P, Giovannetti E, Tiseo M. Resistance mechanisms to osimertinib in EGFR-mutated non-small cell lung cancer. Br J Cancer. 2019;121(9):725–37. Epub 20190930. doi: 10.1038/s41416-019-0573-8. PubMed PMID: 31564718; PMCID: PMC6889286.

20. Enrico D, Tsou FRC, Catani G, Pupareli C, Girotti MR, Alvarez DEU, Waisberg F, Rodriguez A, Reyes R, Chacon M, Reguart N, Martin C. Overcoming Resistance to Osimertinib by T790M Loss and C797S Acquisition Using Gefitinib in a Patient With EGFR-Mutant NSCLC: A Case Report. Jto Clin Res Rep. 2023;4(2). doi: ARTN 100456 10.1016/j.jtocrr.2022.100456. PubMed PMID: WOS:001150075700001.

21. Suzuki M, Uchibori K, Oh-hara T, Nomura Y, Suzuki R, Takemoto A, Araki M, Matsumoto S, Sagae Y, Kukimoto-Niino M, Kawase Y, Shirouzu M, Okuno Y, Nishio M, Fujita N, Katayama R. A macrocyclic kinase inhibitor overcomes triple resistant mutations in EGFR-positive lung cancer. Npj Precis Oncol. 2024;8(1). doi: ARTN 46 10.1038/s41698-024-00542-9. PubMed PMID: WOS:001172604300002.

22. Gunther M, Juchum M, Kelter G, Fiebig H, Laufer S. Lung Cancer: EGFR Inhibitors with Low Nanomolar Activity against a Therapy-Resistant L858R/T790M/C797S Mutant. Angew Chem Int Ed Engl. 2016;55(36):10890–4. Epub 20160728. doi: 10.1002/anie.201603736. PubMed PMID: 27466205.

23. Gunther M, Lategahn J, Juchum M, Doring E, Keul M, Engel J, Tumbrink HL, Rauh D, Laufer S. Trisubstituted Pyridinylimidazoles as Potent Inhibitors of the Clinically Resistant L858R/T790M/C797S EGFR Mutant: Targeting of Both Hydrophobic Regions and the Phosphate Binding Site. J Med Chem. 2017;60(13):5613–37. Epub 20170621. doi: 10.1021/acs.jmedchem.7b00316. PubMed PMID: 28603991.

24. Juchum M, Gunther M, Doring E, Sievers-Engler A, Lammerhofer M, Laufer S. Trisubstituted Imidazoles with a Rigidized Hinge Binding Motif Act As Single Digit nM Inhibitors of Clinically Relevant EGFR L858R/T790M and L858R/T790M/C797S Mutants: An Example of Target Hopping. J Med Chem. 2017;60(11):4636–56. Epub 20170523. doi: 10.1021/acs.jmedchem.7b00178. PubMed PMID: 28482151.

25. Patel H, Pawara R, Ansari A, Surana S. Recent updates on third generation EGFR inhibitors and emergence of fourth generation EGFR inhibitors to combat C797S resistance. Eur J Med Chem. 2017;142:32–47. doi: 10.1016/j.ejmech.2017.05.027. PubMed PMID: WOS:000418208200004.

26. Fukuda S, Suda K, Hamada A, Oiki H, Ohara S, Ito M, Soh J, Mitsudomi T, Tsutani Y. Potential Utility of a 4th-Generation EGFR-TKI and Exploration of Resistance Mechanisms-An In Vitro Study. Biomedicines. 2024;12(7). doi: ARTN 1412 10.3390/biomedicines12071412. PubMed PMID: WOS:001278222100001.

27. Eno MS, Brubaker JD, Campbell JE, De Savi C, Guzi TJ, Williams BD, Wilson D, Wilson K, Brooijmans N, Kim J, Ozen A, Perola E, Hsieh J, Brown V, Fetalvero K, Garner A, Zhang Z, Stevison F, Woessner R, Singh J, Timsit Y, Kinkema C, Medendorp C, Lee C, Albayya F, Zalutskaya A, Schalm S, Dineen TA. Discovery of BLU-945, a Reversible, Potent, and Wild-Type-Sparing Next-Generation EGFR Mutant Inhibitor for Treatment-Resistant Non-Small-Cell Lung Cancer. J Med Chem. 2022;65(14):9662–77. Epub 20220715. doi: 10.1021/acs.jmedchem.2c00704. PubMed PMID: 35838760; PMCID: PMC9340769.

28. Elamin YY, Nagasaka M, Shum E, Bazhenova L, Camidge DR, Cho BC, Felip E, Goto K, Lin CC, Piotrowska Z, Planchard D, Rotow JK, Spigel DR, Tan DSW, Yoshida T, Minchom AR, De Langen A, Kato T, Zalutskaya A, Reckamp KL. BLU-945 monotherapy and in combination with osimertinib (OSI) in previously treated patients with advanced-mutant () NSCLC in the phase 1/2 SYMPHONY study. J Clin Oncol. 2023;41(16). PubMed PMID: WOS:001053772003680.

29. Dingens AS, Haddox HK, Overbaugh J, Bloom JD. Comprehensive Mapping of HIV-1 Escape from a Broadly Neutralizing Antibody. Cell Host Microbe. 2017;21(6):777–+. doi: 10.1016/j.chom.2017.05.003. PubMed PMID: WOS:000403240800016.

30. Schneider WM, Luna JM, Hoffmann HH, Sanchez-Rivera FJ, Leal AA, Ashbrook AW, Le Pen J, Ricardo-Lax I, Michailidis E, Peace A, Stenzel AF, Lowe SW, MacDonald MR, Rice CM, Poirier JT. Genome-Scale Identification of SARS-CoV-2 and Pan-coronavirus Host Factor Networks. Cell. 2021;184(1):120–32 e14. Epub 20201209. doi: 10.1016/j.cell.2020.12.006. PubMed PMID: 33382968; PMCID: PMC7796900.

31. Doud MB, Bloom JD. Accurate Measurement of the Effects of All Amino-Acid Mutations on Influenza Hemagglutinin. Viruses-Basel. 2016;8(6). doi: ARTN 155 10.3390/v8060155. PubMed PMID: WOS:000378848600007.

32. Doud MB, Hensley SE, Bloom JD. Complete mapping of viral escape from neutralizing antibodies. PLoS Pathog. 2017;13(3):e1006271. Epub 20170313. doi: 10.1371/journal.ppat.1006271. PubMed PMID: 28288189; PMCID: PMC5363992.

33. Doud MB, Lee JM, Bloom JD. How single mutations affect viral escape from broad and narrow antibodies to H1 influenza hemagglutinin. Nat Commun. 2018;9. doi: ARTN 1386 10.1038/s41467-018-03665-3. PubMed PMID: WOS:000429689800009.

34. Palacios R, Steinmetz M. Il-3-dependent mouse clones that express B-220 surface antigen, contain Ig genes in germ-line configuration, and generate B lymphocytes in vivo. Cell. 1985;41(3):727–34. doi: 10.1016/s0092-8674(85)80053-2. PubMed PMID: 3924409.

35. Koga T, Suda K, Mitsudomi T. Utility of the Ba/F3 cell system for exploring on-target mechanisms of resistance to targeted therapies for lung cancer. Cancer Sci. 2022;113(3):815–27. Epub 20220123. doi: 10.1111/cas.15263. PubMed PMID: 34997674; PMCID: PMC8898722.

36. Wu NC, Young AP, Al-Mawsawi LQ, Olson CA, Feng J, Qi H, Chen SH, Lu IH, Lin CY, Chin RG, Luan HH, Nguyen N, Nelson SF, Li X, Wu TT, Sun R. High-throughput profiling of influenza A virus hemagglutinin gene at single-nucleotide resolution. Sci Rep. 2014;4:4942. Epub 20140513. doi: 10.1038/srep04942. PubMed PMID: 24820965; PMCID: PMC4018626.

37. Zhang TH, Wu NC, Sun R. A benchmark study on error-correction by read-pairing and tag-clustering in amplicon-based deep sequencing. BMC Genomics. 2016;17:108. Epub 20160212. doi: 10.1186/s12864-016-2388-9. PubMed PMID: 26868371; PMCID: PMC4751728.

38. Damghani T, Wittlinger F, Beyett TS, Eck MJ, Laufer SA, Heppner DE. Structural elements that enable specificity for mutant EGFR kinase domains with next-generation small-molecule inhibitors. Methods Enzymol. 2023;685:171–98. Epub 20230427. doi: 10.1016/bs.mie.2023.03.013. PubMed PMID: 37245901; PMCID: PMC10445336.

39. Vu HA, Xinh PT, Ha HT, Hanh NT, Bach ND, Thao DT, Dat NQ, Trung NS. Spectrum of EGFR gene mutations in Vietnamese patients with non-small cell lung cancer. Asia Pac J Clin Oncol. 2016;12(1):86–90. Epub 20151227. doi: 10.1111/ajco.12448. PubMed PMID: 26707566.

40. Liu Y, Li Y, Ou Q, Wu X, Wang X, Shao YW, Ying J. Acquired EGFR L718V mutation mediates resistance to osimertinib in non-small cell lung cancer but retains sensitivity to afatinib. Lung Cancer. 2018;118:1–5. Epub 20180131. doi: 10.1016/j.lungcan.2018.01.015. PubMed PMID: 29571986.

41. Sharma P, Mahadevia H, Donepudi S, Kujtan L, Gustafson B, Ponvilawan B, Al-Obaidi A, Subramanian J, Bansal D. A Novel EGFR Germline Mutation in Lung Adenocarcinoma: Case Report and Literature Review. Clin Lung Cancer. 2024;25(5):479–82. doi: 10.1016/j.cllc.2024.04.009. PubMed PMID: WOS:001262365600001.

42. Mao L, Zhao W, Li X, Zhang S, Zhou C, Zhou D, Ou X, Xu Y, Tang Y, Ou X, Hu C, Ding X, Luo P, Yu S. Mutation Spectrum of EGFR From 21,324 Chinese Patients With Non-Small Cell Lung Cancer (NSCLC) Successfully Tested by Multiple Methods in a CAP-Accredited Laboratory. Pathol Oncol Res. 2021;27:602726. Epub 20210407. doi: 10.3389/pore.2021.602726. PubMed PMID: 34257561; PMCID: PMC8262202.

43. Si JF, Gu XD, Wang WX, Ying SP, Song ZB. Clinical outcomes of lung adenocarcinoma patients harboring uncommon epidermal growth factor receptor () mutations treated with EGFR-tyrosine kinase inhibitors (TKIs). Ann Palliat Med. 2022;11(5):1624–34. doi: 10.21037/apm-21-2828. PubMed PMID: WOS:000739876000001.

44. Ruiz R, Galvez-Nino M, Santos M, Doimi F, Mas L, Belmar-Lopez C. Mutation pattern of EGFR gene in Peruvian patients with non-small cell lung cancer. J Clin Oncol. 2024;42(16). PubMed PMID: WOS:001275557405288.

45. Ma CH, Liu M, Mu N, Li JD, Li L, Jiang R. Efficacy of afatinib for pulmonary adenocarcinoma with leptomeningeal metastases harboring an epidermal growth factor receptor complex mutation (exon 19del+K754E) A case report. Medicine. 2020;99(43). doi: ARTN e22851 10.1097/MD.0000000000022851. PubMed PMID: WOS:000588186800096.

46. Miyamae Y, Shimizu K, Mitani Y, Araki T, Kawai Y, Baba M, Kakegawa S, Sugano M, Kaira K, Lezhava A, Hayashizaki Y, Yamamoto K, Takeyoshi I. Mutation Detection of Epidermal Growth Factor Receptor and Genes Using the Smart Amplification Process Version 2 from Formalin-Fixed, Paraffin-Embedded Lung Cancer Tissue. J Mol Diagn. 2010;12(2):257–64. doi: 10.2353/jmoldx.2010.090105. PubMed PMID: WOS:000276697300017.

47. Tang HR, Zhang JJ, Hu X, Xu YJ, Dong BG, Wang J, Kong Y, Ma HL, Liao ZX, Zhang JH, Byers LA, Gibbons DL, Glisson BS, Wistuba I, Heymach J, Gomez DR, Futreal A, Chen M. EGFR Mutations in Small Cell Lung Cancer (SCLC): Genetic Heterogeneity and Prognostic Impact. J Thorac Oncol. 2017;12(1):S710–S1. doi: DOI 10.1016/j.jtho.2016.11.936. PubMed PMID: WOS:000413055801436.

48. Robichaux JP, L. X, Vijayan RSK, Hicks JK, Heeke S, Elamin YY, Lin HY, Udagawa H, Skoulidis F, Tran H, Varghese S, He J, Zhang F, Nilsson MB, Hu L, Poteete A, Rinsurongkawong W, Zhang X, Ren C, Liu X, Hong L, Zhang J, Diao L, Madison R, Schrock AB, Saam J, Raymond V, Fang B, Wang J, Ha MJ, Cross JB, Gray JE, Heymach JV. Structure-based classification predicts drug response in EGFR-mutant NSCLC. Nature. 2021;597(7878):732–7. Epub 20210915. doi: 10.1038/s41586-021-03898-1. PubMed PMID: 34526717; PMCID: PMC8481125.

49. Carrera AC, Alexandrov K, Roberts TM. The conserved lysine of the catalytic domain of protein kinases is actively involved in the phosphotransfer reaction and not required for anchoring ATP. Proc Natl Acad Sci U S A. 1993;90(2):442–6. doi: 10.1073/pnas.90.2.442. PubMed PMID: 8421674; PMCID: PMC45679.

50. Rotow JK, Lee JK, Madison RW, Oxnard GR, Jänne PA, Schrock AB. Real-World Genomic Profile of EGFR Second-Site Mutations and Other Osimertinib Resistance Mechanisms and Clinical Landscape of NSCLC Post-Osimertinib. J Thorac Oncol. 2024;19(2):227–39. doi: 10.1016/j.jtho.2023.09.1453. PubMed PMID: WOS:001170894400001.

51. Li M, Qin J, Xie F, Gong L, Han N, Lu H. L718Q/V mutation in exon 18 of EGFR mediates resistance to osimertinib: clinical features and treatment. Discov Oncol. 2022;13(1):72. Epub 20220809. doi: 10.1007/s12672-022-00537-7. PubMed PMID: 35943592; PMCID: PMC9363540.

52. Starrett JH, Guernet AA, Cuomo ME, Poels KE, van Rosenburgh IKV, Nagelberg A, Farnsworth D, Price KS, Khan H, Ashtekar KD, Gaefele M, Ayeni D, Stewart TF, Kuhlmann A, Kaech SM, Unni AM, Homer R, Lockwood WW, Michor F, Goldberg SB, Lemmon MA, Smith PD, Cross DAE, Politi K. Drug Sensitivity and Allele Specificity of First-Line Osimertinib Resistance Mutations. Cancer Res. 2020;80(10):2017–30. doi: 10.1158/0008-5472.Can-19-3819. PubMed PMID: WOS:000535265800017.

53. Morgillo F, Della Corte CM, Fasano M, Ciardiello F. Mechanisms of resistance to EGFR-targeted drugs: lung cancer. Esmo Open. 2016;1(3). doi: UNSP e000060 10.1136/esmoopen-2016-000060. PubMed PMID: WOS:000408015800013.

